# Concentration of intraflagellar transport proteins at the ciliary base is required for proper train injection

**DOI:** 10.1101/2021.08.02.454739

**Authors:** Jamin Jung, Julien Santi-Rocca, Cécile Fort, Jean-Yves Tinevez, Cataldo Schietroma, Philippe Bastin

## Abstract

Construction of cilia and flagella relies on Intraflagellar Transport (IFT). Although IFT proteins can be found in multiple locations in the cell, transport has only been reported along the axoneme. Here, we reveal that IFT concentration at the base of the flagellum of *Trypanosoma brucei* is required for proper assembly of IFT trains. Using live cell imaging at high resolution and direct optical nanoscopy with axially localized detection (DONALD) of fixed trypanosomes, we demonstrate that IFT proteins are localised around the 9 doublet microtubules of the proximal portion of the transition zone, just on top of the transition fibres. Super-resolution microscopy and photobleaching studies reveal that knockdown of the RP2 transition fibre protein results in reduced IFT protein concentration and turnover at the base of the flagellum. This in turn is accompanied by a 4- to 8-fold drop in the frequency of IFT train injection. We propose that the flagellum base provides a unique environment to assemble IFT trains.

## Introduction

Cilia and flagella are constructed by intraflagellar transport (IFT), the movement of protein particles, or IFT trains, along axoneme microtubules (Kozminski et al., 1993; Rosenbaum and Witman, 2002). Trains deliver tubulin and other flagellar precursors to the distal tip of the organelle for incorporation (Craft et al., 2015; Wren et al., 2013). IFT trafficking has been directly visualised using fluorescent proteins in multiple eukaryotic organisms (Orozco et al., 1999)(Engel et al., 2009)(Buisson et al., 2013)(Follit et al., 2006; Williams et al., 2014). Curiously, the largest amounts of IFT proteins are detected in the cytoplasm. In *Chlamydomonas*, cell fractionation revealed that the cell body contains 20 to 50-fold more IFT proteins than the flagellum (Ahmed et al., 2008; Wang et al., 2009). However, neither IFT trafficking nor IFT trains have been reported in the cytoplasm.

IFT proteins are highly concentrated at the base of cilia and flagella (Absalon et al., 2008; Cole et al., 1998; Orozco et al., 1999). In *Chlamydomonas*, immunogold staining with an anti-IFT52 antibody revealed association to the transition fibres, mostly towards the membrane side (Deane et al., 2001). Furthermore, Structured Illumination Microscopy (SIM) showed that IFT proteins make a ring around the flagellar base (Picariello et al., 2019). Similar localisation at the level of the transition fibres was found in *C. elegans* (Jensen et al., 2015; Serwas et al., 2017; Williams et al., 2011) and in RPE-1 cells (Yan et al., 2015). Alterations of the transition fibre proteins CEP164 or DZIP1 resulted in loss of IFT proteins and was accompanied by inhibition of ciliary formation in mammalian cells (Schmidt et al., 2012; Wang et al., 2013; Cajanek & Nigg, 2014). However, this absence precluded further investigation since it was not possible to visualise the impact on transport and IFT train formation itself.

Here, we tried to figure out why IFT concentration at the base could be important for ciliogenesis. We addressed this question in the protist *Trypanosoma brucei* where IFT has been well characterised in live cells (Bertiaux et al., 2018a; Buisson et al., 2013). This organism is amenable to reverse genetics and maintains its existing flagellum whilst assembling the new one, allowing the comparison of mature and growing flagella in the same cell (Sherwin and Gull, 1989). The base of its flagellum displays the typical arrangement with the basal body made of triplet microtubules followed by the transition zone (TZ) composed of doublet microtubules and finally the canonical 9+2 axoneme (Trépout et al., 2017; Vaughan and Gull, 2015). Using a combination of conventional and super-resolution microscopy approaches in both fixed and live cells, we show here that IFT proteins are concentrated as a ring found at the interface of the transition fibres and the proximal portion of the TZ. Following RNAi knockdown of a transition fibre protein, IFT proteins fail to concentrate properly at the base of the flagellum, resulting in a 4- to 8-fold reduction in IFT train injection. These trains traffic more slowly and arrest more frequently. These data suggest that IFT concentration at the level of the transition fibres is necessary for efficient and proper assembly of IFT trains.

## Results and discussion

### IFT proteins form a ring at the level of the transition fibres

To determine the position of IFT proteins at the base of the trypanosome flagellum, double immunofluorescence assays (IFA) were performed with markers of the TZ and the transition fibres. Figure 1A shows the relative position of the IFT basal pool stained with the anti-IFT172 monoclonal antibody (Absalon et al., 2008) compared to the flagellum TZ component FTZC (Bringaud et al., 2000) using widefield microscopy. The merged panel indicates little overlap between the two fluorescent signals and the intensity profile shows that IFT172 can be distinguished from FTZC (Fig. 1A). Next, the relative positioning of the IFT pool with the transition fibres was investigated using the trypanosome retinitis pigmentosa 2 (RP2) protein as marker (Stephan et al., 2007). RP2 can be detected with the monoclonal antibody YL1/2 (Andre et al., 2014; Stephan et al., 2007) or following *in situ* expression of a fusion protein with mCherry (mCh, Fig. S1). The IFT172 pool can be distinguished from the RP2 signal but seems to overlap partially as seen from the intensity profile (Fig 1B, right panel).

**Figure 1.**
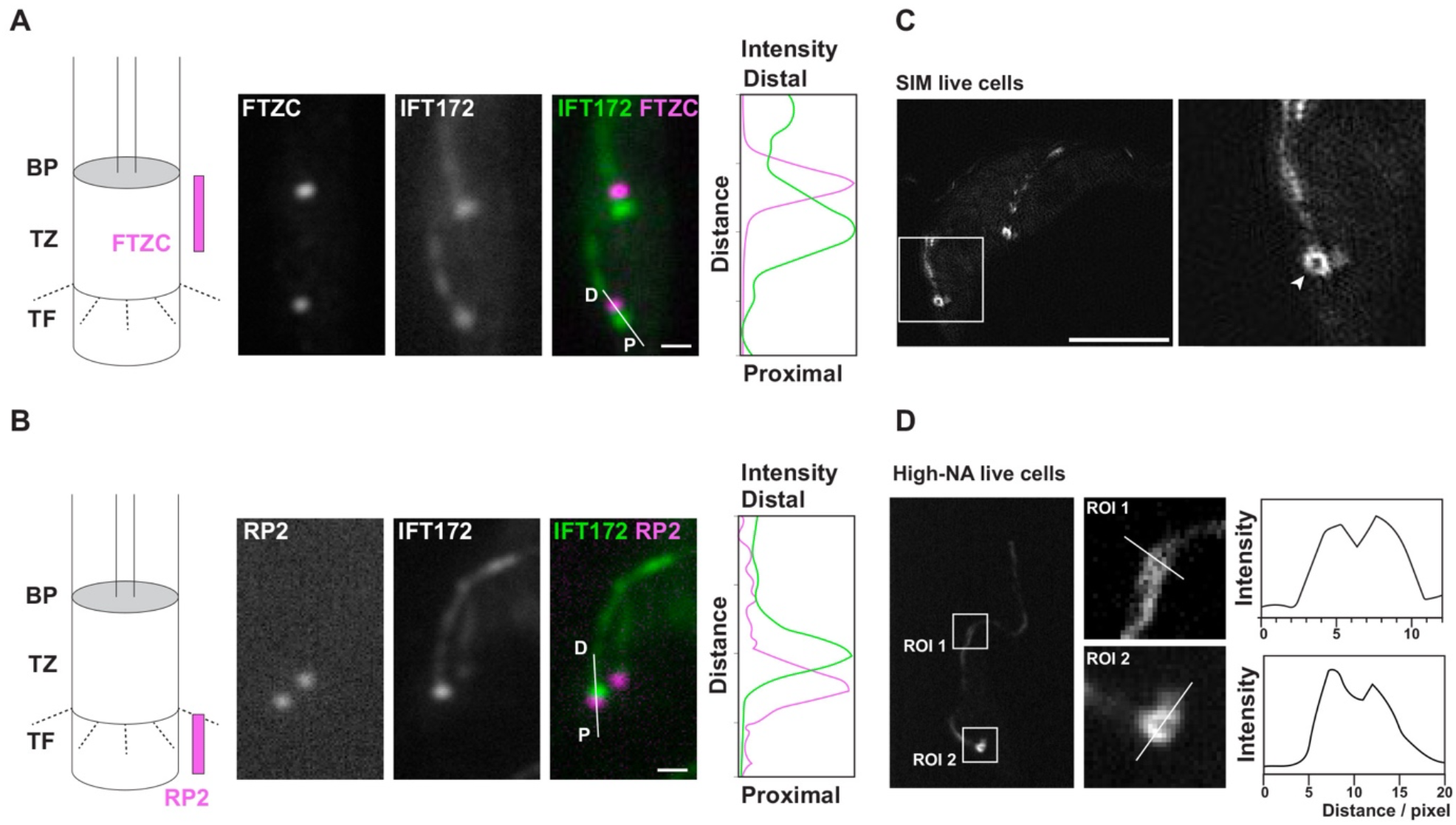
The IFT pool is localized at the proximal TZ in direct vicinity to the transitional fibre marker RP2. Trypanosomes expressing mCh::RP2 were fixed using the PFA-methanol protocol and processed for IFA with various markers revealing that the IFT pool localizes in between the distal transition zone marker FTZC and the transitional fibre marker RP2. **(A)** The IFT pool was stained with the anti-IFT172 antibody and the TZ with FTZC as indicated. The intensity profile was measured along the proximal (P) and distal (D) axis and shows the respective localization of IFT172 (green) and FTZC (magenta). **(B)** Staining of IFT172 (green) and RP2 (magenta) and corresponding plot profile. The IFT pool can be clearly distinguished from the RP2 signal but overlaps partially as shown on the profile plot. **(C)** Structured Illumination Microscopy (SIM) of a biflagellated live cell expressing GFP::IFT52 revealing the ring like structure of the IFT pool (arrowhead on magnified panel). **(D)** High-NA live imaging of a trypanosome expressing mNG::IFT81. Regions of interest along the flagellum (ROI1) or its base (ROI2) are magnified on the left. Two parallel tracks of IFT signal within the axoneme are detected, in agreement with the known IFT train distribution. The IFT signal at the proximal end of the flagellum appears as a ring. Cartoons in (A-B) indicate the organisation of the base of the flagellum with the transition fibres (TF), the transition zone (TZ) and the basal plate (BP). The position of known markers is shown in magenta. Scale bars=2.5 µm (A-B), 5 µm (C).

Given the complexity of this region, we turned towards super-resolution microscopy using direct Optical Nanoscopy with Axial Localized Detection (DONALD)(Bourg et al., 2015). *T. brucei* cells expressing mCh::RP2 were used and dual IFA was performed with the monoclonal antibody against IFT172 and with an antibody against mCherry. The cell presented at Figure 2A possesses two flagella. In the proximal region of the growing flagellum, two parallel lines are observed for IFT172, presumably corresponding to the two sets of microtubule doublets of the axoneme where IFT trains are observed in trypanosomes (Bertiaux et al., 2018a). The higher concentration of IFT172 at the flagellum base is easily visible. The signal partially overlaps with RP2 but is found in a more apical position (Fig. 2A). Figure 2B shows a representative side view image of the TZ region indicating that the IFT pool (green) is distal of the transitional fibres (magenta) with a region of overlap. The IFT pool is therefore on top of the transition fibres and present in the proximal portion of the TZ. In addition, IFT172 signal is also detected as thin lines along the distal portion of the TZ, possibly representing IFT trains under construction (Fig. 2B). These images are compatible with the progression of IFT material observed during train construction in *C. elegans* (Prevo et al., 2015) and RPE-1 cells (Yang et al., 2019).

**Figure 2.**
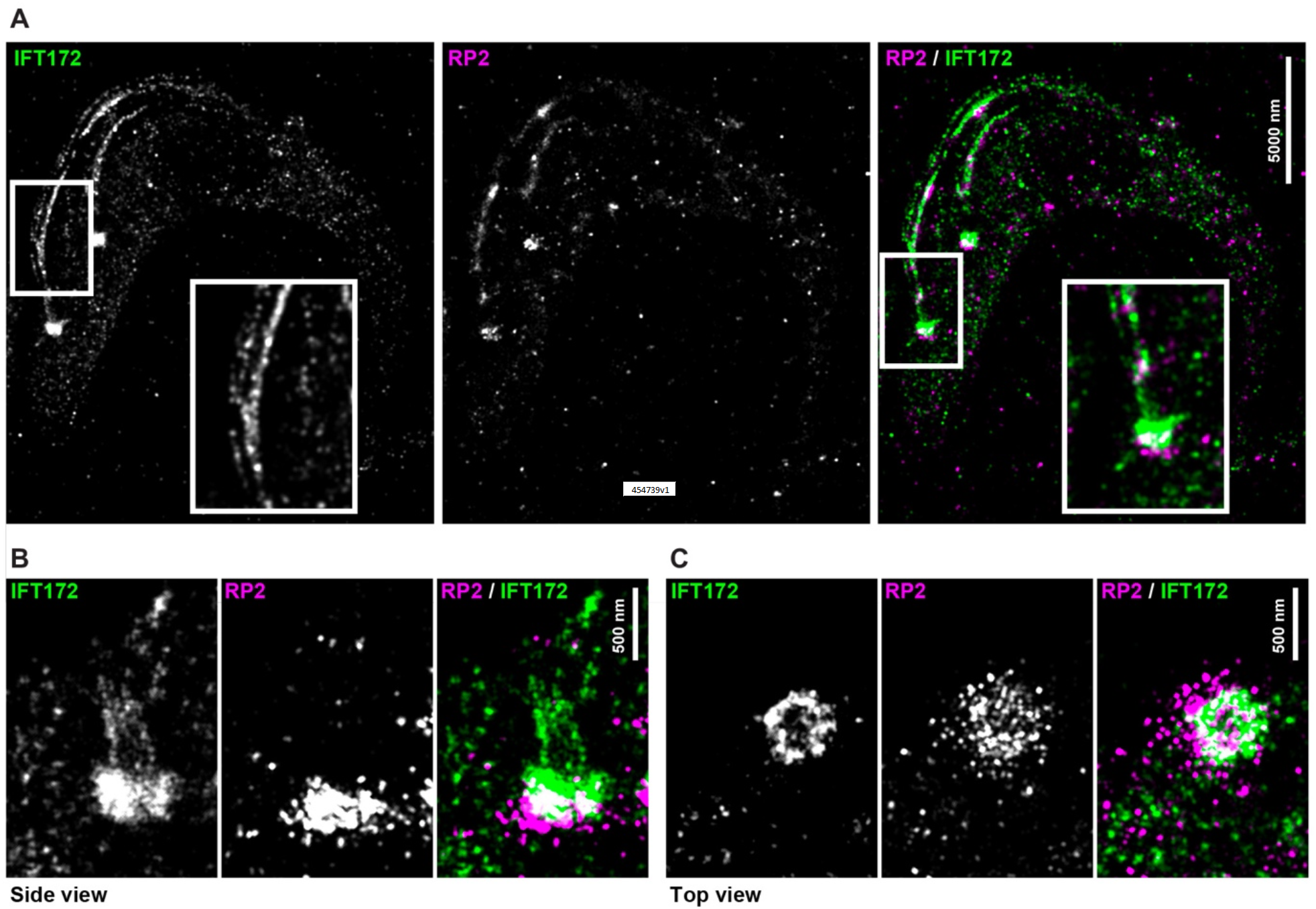
Two-colour 3D-STORM confirms the ring structure of the IFT pool and its relative position to the transition fibres. The position of IFT172 was investigated by direct optical nanoscopy with axially localized detection (Bourg et al., 2015) on fixed *T. brucei* cells expressing mCh::RP2 using antibodies against IFT172 (green) and mCherry (magenta) as indicated. **(A)** View of a whole cell with two flagella. The inset on the left panel is a zoom on the proximal region of the new flagellum showing IFT train distribution on both sides of the axoneme. The inset on the right panel is a magnification of the base of the flagellum where IFT172 and mCh::RP2 are in close proximity. **(B)** Representative side view image of the TZ region indicating that the IFT pool is distal of the transition fibre protein RP2. The merge in white shows the partial overlap. **(C)** Image of a representative top view of the TZ region. IFT172 is arranged in a ring like structure. The scale bars are 500 nm.

A representative top view of the base of the flagellum reveals that IFT172 is arranged in a ring like structure (Fig. 2C). This pattern was confirmed in 16 analysed 3D STORM stacks. The diameter of the IFT172 ring was measured at 356 ± 26 nm (n=16), which is very close to the diameter of the flagellar membrane that surrounds the TZ as determined by scanning transmission electron microscopy (STEM)(352 ± 27 nm)(Trépout et al., 2017). This is compatible with the presence of the IFT pool as a ring around the 9 doublet microtubules at the proximal portion of the TZ, overlapping with the transition fibres.

Since fixation could alter the distribution of IFT proteins that are soluble and highly dynamic, we investigated their location in live cells by two approaches. First, cells expressing a GFP::IFT52 fusion protein (Absalon et al., 2008) were observed by Structured Illumination Microscopy (SIM). Two lines were visible along the axoneme, in agreement with the restriction of IFT trains along doublets 3-4 and 7-8 (Bertiaux et al., 2018a). IFT proteins were detected as a ring at the base of the flagellum (Fig. 1C), confirming the STORM data was not an artefact due to fixation. Since GFP::IFT52 is overexpressed in that cell line, we reproduced the observation with trypanosomes expressing an endogenously mNeonGreen tagged IFT81 using high-resolution microscopy (Fig. 1D) with an objective with a 1.57 numerical aperture (Bertiaux et al., 2018a).

This combination of approaches with live and fixed cells demonstrates that the basal pool of IFT proteins is positioned on top of the transition fibres, forming a ring-like structure surrounding the 9 doublet microtubules.

### Knockdown of a transition fibre component affects IFT concentration at the base of the flagellum

We reasoned that affecting the composition of the transition fibres could impact IFT distribution and help understanding the significance of this concentration. RP2 was the only transition fibre protein characterised in trypanosomes (Stephan et al., 2007) when this study was initiated. Tetracycline-inducible RNAi knockdown (Wang et al., 2000) of RP2 was performed in the cell line expressing mCh::RP2 fusion as a reporter. Quantification of the mCh::RP2 signal at the base of the flagellum revealed the loss of the signal and validated the efficiency of RNAi knockdown (Fig. S1A). Furthermore, IFA with the monoclonal antibody YL1/2 that detects both tyrosinated tubulin and RP2 (Andre et al., 2014) confirmed the efficiency of RP2 knockdown (Fig. S1B). This impacted on both cell growth (Fig S2A) and flagellum length (Fig S2B), as expected (Stephan et al., 2007).

The impact of RP2 depletion on the abundance of IFT proteins was evaluated by western blotting with the anti-IFT172 monoclonal antibody (Fig. S3A). The total amount was not visibly affected, showing that RP2 knockdown did not interfere with IFT protein expression and turnover. By contrast, IFA revealed that IFT172 concentration along the length of the flagellum was significantly decreased (Fig. S3B). Furthermore, the concentration of IFT at the base was reduced when stained with the anti-IFT172 in fixed cells (Fig. S4A). This was not a consequence of fixation as this result was reproduced with live *RP2*^*RNAi*^ cells expressing YFP::IFT81 as IFT reporter (Fig. S4B).

Given the limitations of conventional microscopy, 3D-STORM was used to analyse IFT distribution at the base of the flagellum, comparing control cells and cells in which RP2 was knocked-down by RNAi for 2 days. Representative side view images show that in control cells, the IFT pool displayed the expected concentration at the base of the flagellum (Fig. 3A, left panel). By contrast, the distribution pattern of IFT172 is altered in cells depleted of RP2 and the signal looks spread throughout the TZ region (Fig. 3A, right panel). Semi-automated measurements on representative top views were carried out in order to measure the IFT pool diameter. In contrast to control cells, the algorithm was not able to identify any specific structures in the RP2 knockdown situation. This indicates severe perturbation of the IFT concentration at the flagellum base.

**Figure 3.**
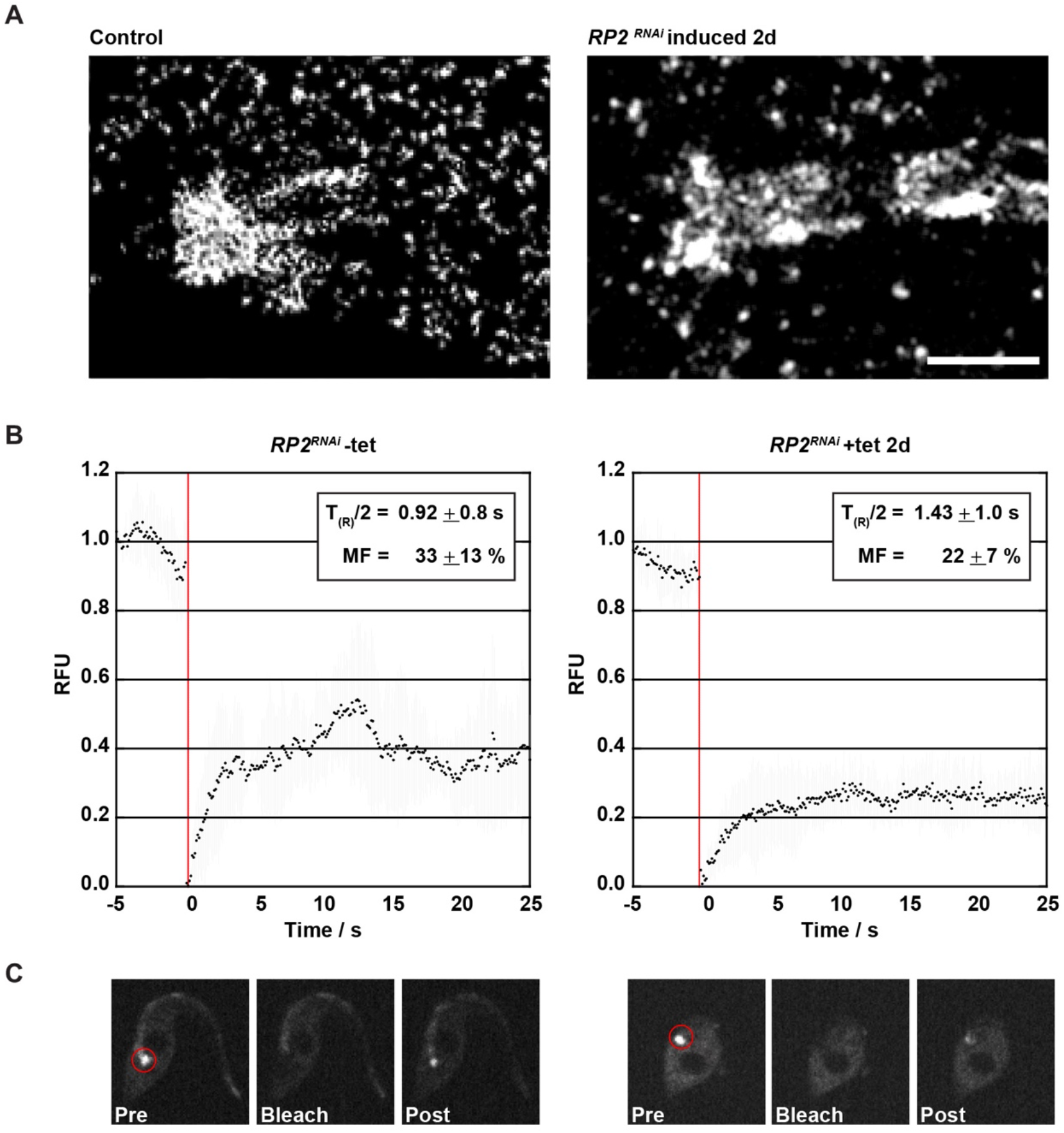
RNAi knockdown of RP2 leads to the redistribution of IFT172 and reduced IFT dynamics. **(A)** 3D-STORM was used to analyse IFT distribution comparing control cells and cells in which RP2 was depleted by RNAi for 2 days. **(A)** Representative side view images show that the IFT pool in control cells displays the expected concentration at the proximal TZ region whereas in *RP2*^*RNAi*^ cells induced for 2 days (d), the distribution pattern of IFT is altered. Scale bar=500 nm. **(B-C)** Fluorescence recovery analysis following bleaching of the IFT pool at the base of the flagellum (red circle in C) in *RP2*^*RNAi*^ cells expressing mNG::IFT81. Control cells (without RNAi induction, Video S1) and cells induced for 2 days (Video S2) were studied in parallel. **(B)** Quantification in relative fluorescence units (RFU) of the signal in the region of interest. Data were recorded 5 seconds before bleaching and 25 seconds after. The mean fluorescence values with standard deviation are plotted for at least 10 cells per condition. **(C)** Representative images with the photobleached region (red circle) before (Pre), immediately after the bleach (Bleach) and at the recovery plateau (post).

Next, we asked whether this modification could affect IFT dynamics at the base of the flagellum. First, we photo-bleached the IFT pool at the base and monitored its recovery (FRAP) in cells endogenously expressing mNeonGreen::IFT81 as reporter of IFT in the *RP2*^*RNAi*^ cell line (Fig. 3B). Cells were analysed in the absence of RNAi induction or in RP2 knockdown conditions and recovery of the IFT81 fluorescence at the flagellum base was recorded (Fig. 3B, Video S1). In control conditions, the signal recovered rapidly, consistently with the recycling of fluorescent IFT proteins present in trains in the flagellum before bleaching, reaching a plateau at ∼33%, comparable with published data (Buisson et al., 2013; Hibbard et al., 2021). The picture turned out to be different in RP2 knockdown cells where two differences in the IFT dynamics were detected (Fig. 3B, right panel & Video S2). First, the recovery half-life is prolonged after RP2 knockdown and second, the mobile fraction of the tagged IFT is reduced from 33 to 22% (Fig. 3B). This shows that IFT dynamics are reduced (but not abolished) after RP2 knockdown. We conclude that depletion of the transition fibre RP2 protein interferes with the concentration, dynamics and proper distribution of IFT proteins at the base of the flagellum.

### IFT injection and trafficking relies on IFT concentration at the base

To evaluate the impact of the modified distribution of IFT proteins at the base of the flagellum on IFT trafficking, live cell imaging and kymograph analyses of *RP2*^*RNAi*^ cells expressing mNG::IFT81 were performed. In non-induced *RP2*^*RNAi*^ cells, robust IFT was detected, with sustained injection of anterograde trains (Fig. 4A, left panel & Video S3), as previously reported (Buisson et al., 2013). After 24h of RNAi knockdown, three major modifications of IFT trafficking were observed: the number of injected trains appeared lower, their progression in the flagellum looked more chaotic and apparently immotile patches of IFT signals were observed at various positions along the flagellum (Video S4). These phenotypes were more exacerbated after 48h of RNAi knockdown, with sometimes large amounts of mNG::IFT81 signal stuck on the flagellum and much reduced IFT trafficking (Fig. 4A, right panel & Video S5).

**Figure 4.**
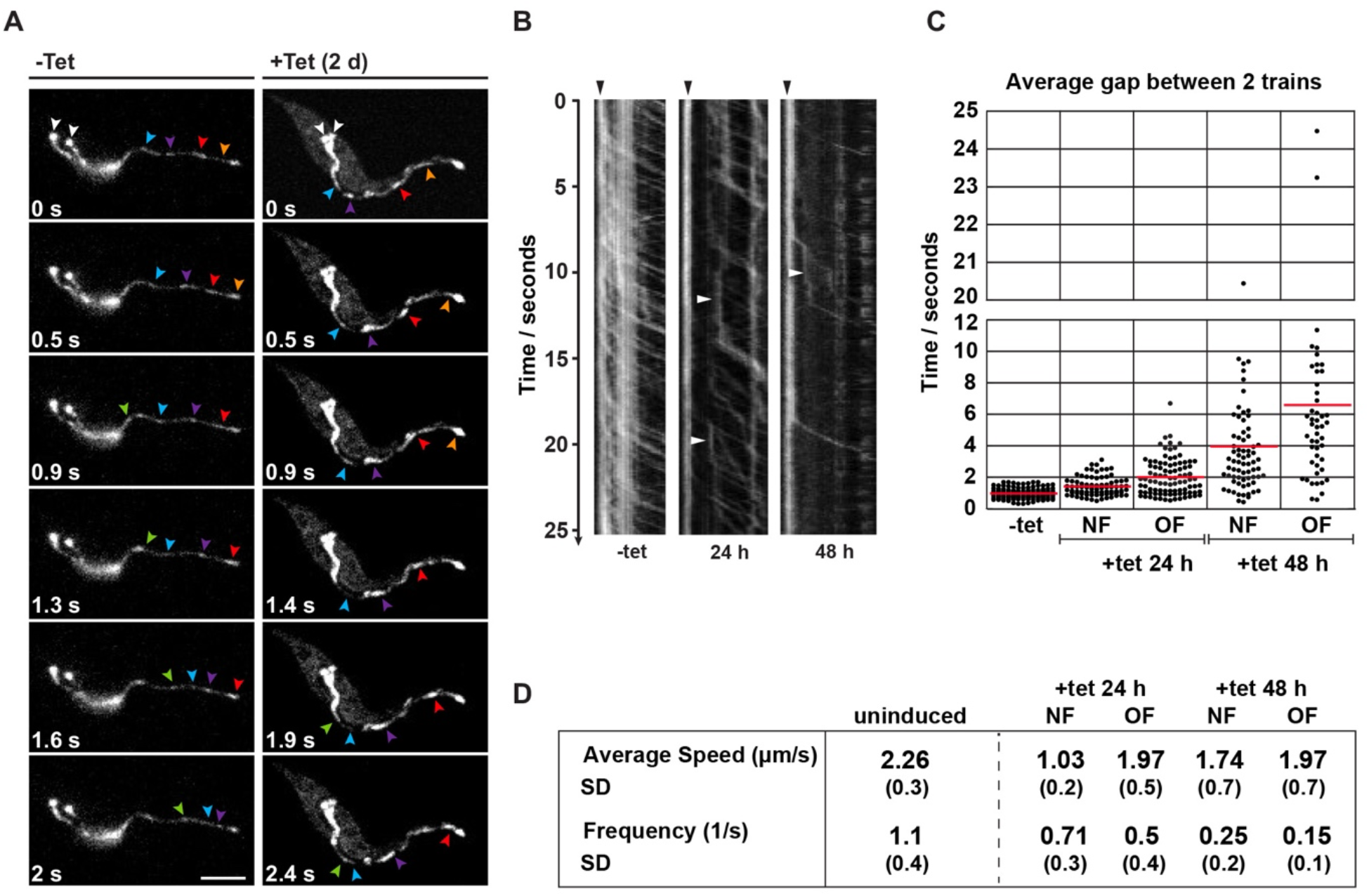
RNAi knockdown of RP2 alters IFT trafficking. *RP2*^*RNAi*^ cells expressing mNG::IFT81 were grown in absence or presence of tetracycline (tet) and IFT was recorded for 25-30 seconds. **(A)** The flagellum base is indicated with white arrowheads and coloured arrowheads point at various IFT trains at the indicated time of recording. See corresponding Videos 3-5. **(B)** Kymograph analysis was performed in cells in the indicated conditions. Robust IFT is detected in non-induced conditions but trains become less frequent during the course of RNAi knockdown. Black and white arrowheads indicate the IFT basal pool and standing IFT trains, respectively. Note that some of these trains can resume motility. **(C-D)** Quantification of the indicated parameters in control (n=180 trains) and knockdown (n=539 trains) conditions. NF = new flagellum, OF = old flagellum.

IFT trafficking was quantified using kymograph analysis (Fig. 4B). In control non-induced conditions, trains were injected at a frequency of 1.1 per second and travelled at an average speed of 2.26 µm per second (Fig. 4D), as previously observed (Buisson et al., 2013). IFT was quantified in growing and mature flagella of knockdown cells, revealing reduced IFT train frequency and velocity in both (Fig. 4C,D). After 2 days in RNAi conditions, the frequency of injection became erratic, falling at 0.15 train per second in the old flagellum, i.e. a reduction of 8-fold compared to the control situation. This is better illustrated as the time gap separating the injection of two anterograde trains that increases to an average 5 s and reaches up to 25 s (Fig. 4C). Another striking observation was an increase in the number of arrested trains (Fig. 4B, arrowheads) corresponding to the immotile patches of IFT signal observed on live cells (Videos S4 and S5). Kymograph analysis revealed that these trains did not arrest for ever since many of them were seen moving again after several seconds (Fig. 4B, arrowheads).

We conclude that depletion of RP2 leads to a reduced concentration of IFT proteins at the flagellar base and this impacts on both injection and trafficking of IFT trains. This suggests that proper assembly of IFT trains requires the local high concentration of IFT proteins and motors at the transition fibres, coupled to the proximity of membrane and microtubule doublets. This combination of parameters is only observed in that location and could explain why train formation and movement is restricted to cilia and flagella, despite the presence of IFT proteins in other locations. The reduced concentration of IFT material at the base of the flagellum in the absence of RP2 could be explained by direct interactions between IFT proteins and transition fibre proteins (Wei et al., 2013) or if the peculiar environment of the TZ allows retention of IFT proteins (Hibbard et al., 2021). In agreement with this proposal, knockdown of RP2 affects the localisation of MKS1 and MKS6 proteins in trypanosomes (Andre et al., 2014), suggesting perturbation of the local environment.

It was initially proposed that the construction of shorter flagella in *RP2*^*RNAi*^ cells was explained by a role for RP2 in processing tubulin prior to its entry in the flagellum (Stephan et al., 2007). However, further analyses questioned that interpretation (Andre et al., 2014). We suggest that reduced IFT train frequency is responsible for the assembly of shorter flagella, as previously observed when the IFT kinesin motor was depleted in trypanosomes (Bertiaux et al., 2018b) or in a temperature-sensitive mutant of *Chlamydomonas* (Marshall and Rosenbaum, 2001).

## Materials and Methods

### Trypanosome cell lines and cultures

Cell lines used for this work are derivatives of *T. brucei* strain 427 cultured at 27°C in SDM79 medium supplemented with hemin and 10% foetal calf serum (Brun and Schonenberger, 1979). For N-terminal tagging of RP2 with mCherry, the first 497 nucleotides of *RP2* (Tb927.10.14010) were chemically synthesised by GeneCust Europe for in-frame cloning using the Hind III and ApaI restriction sites of the p2845 vector (Kelly et al., 2007). The plasmid was linearised with Eco811 for integration in the *RP2* locus, hence ensuring *in situ* expression under the control of the *RP2* 3’ untranslated region. For *RP2* RNAi knockdown, the same sequence was cloned in the pZJM vector (Wang et al., 2000) using HindIII and XhoI sites. The pZJM plasmid contains two tetracycline-inducible T7 promoters facing each other, ensuring inducible expression of *RP2* double-stranded RNA when transformed in the 29-13 cell line that constitutively expresses the T7 RNA polymerase and the tet-repressor (Wirtz et al., 1999). For imaging IFT, the *RP2*^*RNAi*^ cell line was transformed with the previously reported vector p2675mNGIFT81 (Bertiaux et al., 2018a), p2675YFPIFT81 (Bhogaraju et al., 2013) or pPCPFRGFPDHC2.1 (Blisnick et al., 2014). Linearised plasmids were nucleofected (Burkard et al., 2007), resistant cells were grown in media with the appropriate antibiotic concentration and clonal populations were obtained by limited dilution. The cell line expressing GFP::IFT52 has been described previously (Absalon et al., 2008).

### Immunofluorescence imaging

Trypanosomes were fixed either in methanol for 5 minutes at -20°C (Sherwin and Gull, 1989) or using a sequential paraformaldehyde-methanol fixation (Bertiaux et al., 2018a). Briefly, cultured parasites were washed twice in SDM79 medium without serum and spread directly onto poly-L-lysine coated slides (Fisher Scientific J2800AMMZ). Cells were left for 10 minutes to settle prior to treatment with 1 volume 4% PFA solution in PBS at pH 7. After 5 minutes, slides were washed briefly in PBS before being fixed in pure methanol at a temperature of -20°C for 5 minutes followed by a rehydration step in PBS for 15 minutes. For immunodetection, slides were incubated for 1 h at 37°C with the appropriate dilution of the first antibody in 0.1% BSA in PBS; a monoclonal antibody against the IFT-B protein IFT172 (Absalon et al., 2008), a polyclonal antiserum against FTZC (Bringaud et al., 2000), a monoclonal antibody against the TbSAXO1 protein as marker of the axoneme (Dacheux et al., 2012), the YL1/2 rat monoclonal antibody that detects both tyrosinated tubulin and RP2 (Andre et al., 2014; Kilmartin et al., 1982) or a commercial antibody against dsRed that detects mCherry (Living Colors DsRed Polyclonal Antibody, Clontech). After 3 consecutive 5-min washes in PBS, species and subclass-specific secondary antibodies coupled to the appropriate fluorochrome (Alexa 488, Cy3, Jackson ImmunoResearch) were diluted 1/400 in PBS containing 0.1% BSA and were applied for 1 h at 37°C. After washing in PBS as indicated above, cells were stained with a 1µg/ml solution of the DNA-dye DAPI (Roche) and mounted with the Slowfade antifade reagent (Invitrogen). Slides were immediately observed with a DMI4000 microscope (Leica) with a 100X objective (NA 1.4) using a Hamamatsu ORCA-03G camera with an EL6000 (Leica) as light excitation source. Image acquisition was performed using Micro-manager software and images were analysed using ImageJ (National Institutes of Health, Bethesda, MD).

### Structured illumination microscopy (SIM)

Live-cell SIM super-resolution imaging with SIM was performed as described in (Bertiaux et al., 2018a) during a visit at the Advanced Imaging Centre of the Janelia Farm campus. The cell line expressing GFP::IFT52 and TdT::IFT81 was grown in standard conditions, and samples were mounted between glass and coverslip for observation on a custom-built microscope (Gustafsson, 2000; Gustafsson et al., 2008) based on an AxioObserver D1 stand equipped with an UAPON100XOTIRF 1.49 NA objective (Olympus) and an Orca Flash 4.0 sCMOS camera (Hamamatsu Photonics). GFP fluorophores were excited with a 488-nm laser (500 mW; SAPPHIRE 488–500; Coherent) and detected through an adequate emission filter (BP 500–550 nm). The sequence contains a series of 300 images exposed for 20 ms each for a total duration of 17.7 s.

### High-resolution imaging of IFT trafficking

Cell lines were grown in standard conditions and samples were mounted between glass and quartz coverslip (Cover glasses HI, Carl Zeiss, 1787-996). For movie acquisition, a spinning disk confocal microscope (UltraVIEW VOX, Perkin-Elmer) equipped with an oil immersion objective Plan-Apochromat 100x/1.57 Oil-HI DIC (Carl Zeiss) was used. Movies were acquired using Volocity software with an EMCCD camera (C-9100, Hamamatsu) operating in streaming mode. The samples were kept at 27°C using a temperature-controlled chamber. Sequences of 30 seconds were acquired with an exposure time of 100 milliseconds per frame. Kymographs were extracted and analysed with Icy software (de Chaumont et al., 2012) using the plug-in Kymograph Tracker 2 (Chenouard et al., 2010).

### Fluorescence recovery after photobleaching (FRAP)

For FRAP analysis of induced or non-induced *RP2*^*RNAi*^ cells expressing mNG::IFT81, a Zeiss inverted microscope (Axiovert 200) equipped with an oil immersion objective (magnification x63 with a 1.4 numerical aperture) and a spinning disk confocal head (CSU22, Yokogawa) was used (Buisson et al., 2013). Images were acquired using Volocity software with an EMCCD camera (C-9100, Hamamatsu) operating in streaming mode. A sample was taken directly from the culture grown at 6-8 × 10^6^ cells/mL and trapped between slide and coverslip. The samples were kept at 27°C using a microscope incubation chamber. Time-lapse sequences were acquired to analyse signal recovery after photobleaching. Movies were taken using a time lapse of 2 minutes. Exposure time was 0.1 second per frame (binning was 1×1 pixels). In this case, 5 cells were identified per series and their position recorded. IFT was monitored for 20 seconds before photobleaching. After bleaching of the IFT pool at the flagellar base, sequences of at least 100 seconds were filmed for each with an exposure time of 0.1 second per frame in order to record fluorescence recovery.

### Super-resolution imaging by direct optical nanoscopy with axially localized detection (3D-STORM)

Wild-type trypanosomes, cells expressing mCh::RP2 or *RP2*^*RNAi*^ cells induced for 2 days were fixed and labelled for dSTORM imaging using antibodies against IFT172, revealed with an Alexa Fluor 647-coupled secondary antibody against mouse IgG1 (Molecular Probes A21237, 1/500), and antibodies against dsRed that detect mCherry, revealed with an Alexa Fluor 555-coupled secondary antibody against rabbit antibodies (Molecular Probes A27039, 1/500). 2×10^7^ cells per round 25 mm coverslip were harvested and fixed using the sequential PFA methanol protocol described above. Primary and secondary antibodies were incubated for 45 minutes at 37°C in a humid environment. After the final wash cells were post-fixed with 4% PFA for 5 minutes followed by two additional washes in PBS. Samples were stored in PBS until acquisition. For dual colour imaging, a sequential strategy was adopted, imaging first the AF647-labelled proteins and afterwards the AF555-labelled proteins. Labelled cells were incubated in dSTORM buffer (SMART kit, Abbelight), and immediately imaged an Abbelight SAFe360 module in DONALD configuration (Bourg et al., 2015; Deschamps et al., 2014). This enables the detection of single fluorophores with a lateral localization precision of typically 5 to 10 nm and an absolute vertical position with a precision of 15 nm and it is optimally adapted for 3D super resolution imaging of biological structures within 500nm of the coverslip. The dual view super resolution module SAFe360 was equipped with two Hamamatsu Orca FLASH4 sCMOS cameras and mounted on an Olympus IX83 inverted microscope with a 1.49NA 100X oil objective and ZDC focus control; the sample was excited in HiLO using a 640nm laser (for AF647) and a 532nm laser (for AF555), with a Starscan azimuthal TIRF (Errol, France). Once established optimal photo-switching, in DONALD configuration two simultaneous acquisition films were collected (50 ms per frame, 50000 frames per acquisition) and used to perform lateral and axial localisation of individual fluorophores with Abbelight NEO software. Drift correction, data post-processing and visualization were also performed using NEO software.

### Western blot

Cells were washed once in Phosphate Buffer Saline (PBS). Laemmli loading buffer was added to the cells and samples were boiled for 5 minutes. 20μg of protein were loaded into each lane of a Criterion™ XT Bis-Tris Precast Gel 4-12% (Bio-Rad, UK) for SDS-Page separation. XT-Mops (1X) diluted in deionised water was used as a running buffer. Proteins were transferred onto nitrocellulose membranes using the BioRad^®^ Trans-Blot Turbo™ blotting system (25V over 7 minutes). The membrane was blocked with 5% skimmed milk overnight and then incubated with the monoclonal anti-IFT172 antibody (Absalon et al., 2008) diluted 1/500 in 0.1% milk and 0.1% Tween in PBS (PBST). As a loading control the anti-PFR L13D6 (Kohl et al., 1999) and 2E10 (Ismach et al., 1989) monoclonal antibodies were used. After primary antibody incubation, three washes of 5 minutes each were performed in 0.05% PBST followed by secondary antibody incubation. Anti-mouse secondary antibody coupled to horseradish peroxidase, diluted to 1/20,000 in 0.05% PBST containing 0.1% milk, and the membrane was incubated with this for 1 hour. The Amersham ECL Western Blotting Detection Reagent Kit (GE Healthcare Life Sciences, UK) was used for final detection of proteins on the membrane.

### Statistical analyses

In absence of other indications, all errors correspond to the standard deviation of the population. Anova tests were performed using the appropriate tool in Kaleidagraph v4.5.2. Comparison of the mean of cell density in culture growth experiments was done at each time point between the control and the test conditions using Student’s T test, adjusted for unequal variances when necessary (Prisma GraphPad v8). Using the same test, flagellum length and protein fluorescence were compared at each time point to the pre-induction condition (D0) with the same cell cultures. In the graphs, * stands for p ≤ 0.05; ** for p ≤ 0.01; and *** for p ≤ 0.001 as compared to the control.

## Acknowledgements

We thank Derrick Robinson and Frédéric Bringaud (Bordeaux University, France) for providing the mAb25 and anti-FTZC antibodies, respectively. We are grateful to the Photonic Bioimaging and Ultrastructural Bioimaging facilities for providing access to their equipment. SIM was performed at the Advanced Imaging Centre, Janelia Research Campus, jointly sponsored by the Howard Hughes Medical Institute and the Gordon & Betty Moore Foundation. We are grateful to Jim Morris for having made this project feasible and to Lin Shao for training and advice. We thank Aline Alves, Serge Bonnefoy and Thierry Blisnick for critical reading of the manuscript.

J.S-R. was funded by post-doctoral fellowships from la Fondation pour la Recherche Médicale (SPF20120523874) and by the Institut Carnot Pasteur-Maladies Infectieuses. C. F. was supported by fellowships from French National Ministry for Research and Technology (doctoral school CDV515) and from La Fondation pour la Recherche Médicale (FDT20150532023). This work was funded by an ANR grant (14-CE35-0009-01), by La Fondation pour la Recherche Médicale (Equipe FRM DEQ20150734356) and by a French Government Investissement d’Avenir programme, Laboratoire d’Excellence “Integrative Biology of Emerging Infectious Diseases” (ANR-10-LABX-62-IBEID). Live imaging at the AIC, Janelia Farm Research Campus was supported by the Citech (Centre d’Innovation et Recherche Technologique) of the Institut Pasteur.

## Author contributions

J. Santi-Rocca and J. Jung conceptualised the study, performed experiments and contributed to analysis. C. Fort, J.Y. Tinevez and C. Schietroma performed experiments and contributed to analysis. P. Bastin conceived and coordinated the project and wrote the paper with contributions from all authors.

The authors declare no competing financial interests.

## Legends for supplementary material

**Figure S1.**
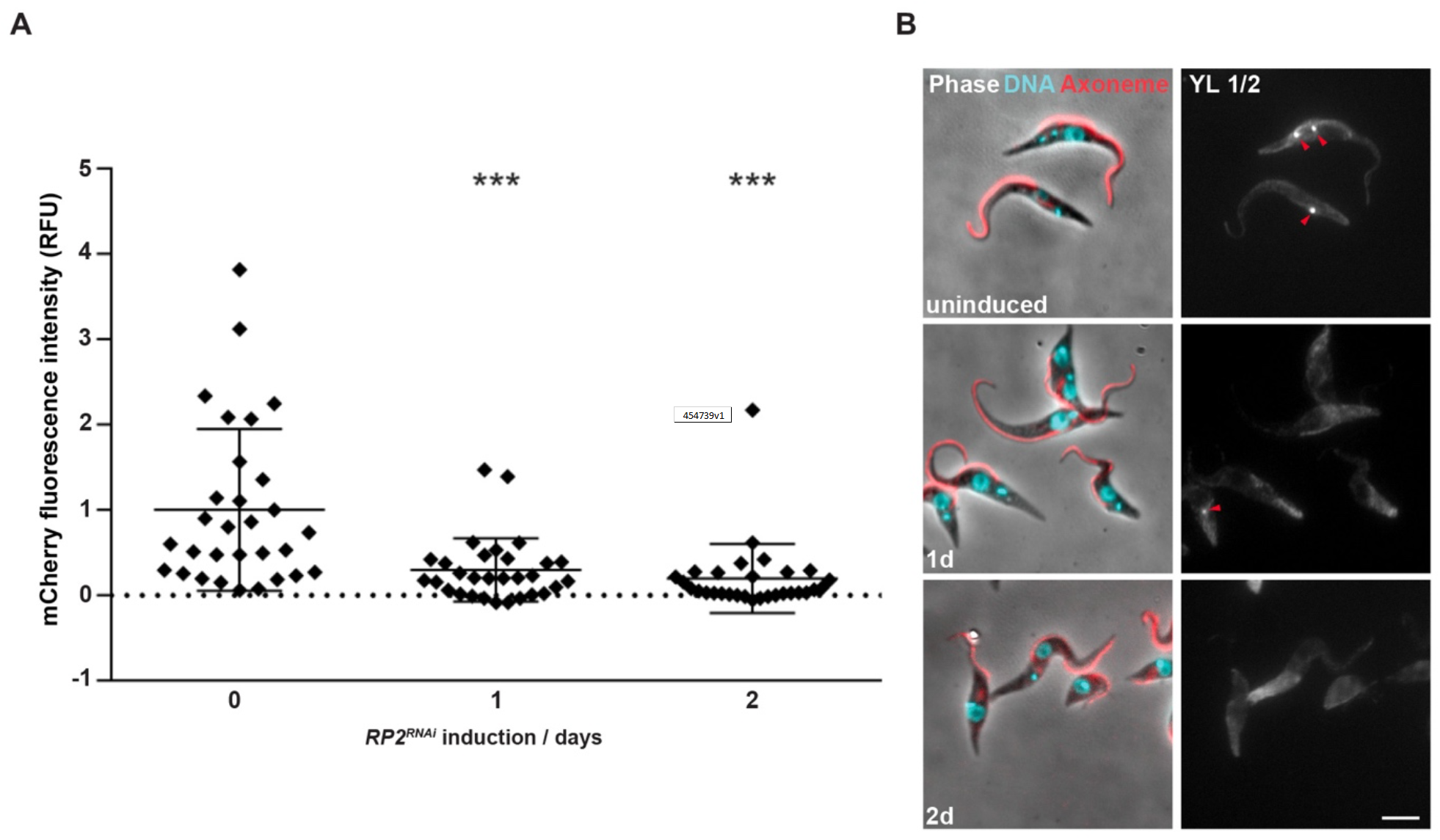
Efficiency of RP2 knockdown. **(A)** The efficiency of RP2 protein depletion was evaluated in the *RP2*^*RNAi*^ cell line expressing an mCh::RP2 fusion protein from its endogenous locus and a YFP::IFT81 reporter. The fluorescence intensity of the mCherry at the base of the flagellum was measured using mCherry direct fluorescence in uniflagellated live cells (n=30, 10 parasites from 3 different experiments) grown without induction or after one or two days of culture in the presence of tetracycline to induce expression of *RP2* dsRNA. The signal becomes virtually null after 2 days in RNAi conditions. * stands for p ≤ 0.05; ** for p ≤ 0.01; and *** for p ≤ 0.001 as compared to the control **(B)** IFA experiment carried out on fixed *RP2*^*RNAi*^ cells in the indicated conditions of RNAi induction using the mAb25 antibody as a marker of the axoneme (red, merged with the phase contrast image and with DAPI staining in blue, left panel) and YL1/2 that detects both RP2 and tyrosinated tubulin (white, right panel)(Andre et al., 2014). The arrowheads indicate the base of the flagellum where RP2 can be detected. While all the cells possess RP2 at the flagellum base without induction, only a few cells remain positive at that location after one day (1d) of RNAi and the vast majority becomes negative after two days (2d). Nevertheless, staining of tyrosinated tubulin in the corset and on the growing flagellum is unchanged. Scale bar: 5 µm.

**Figure S2.**
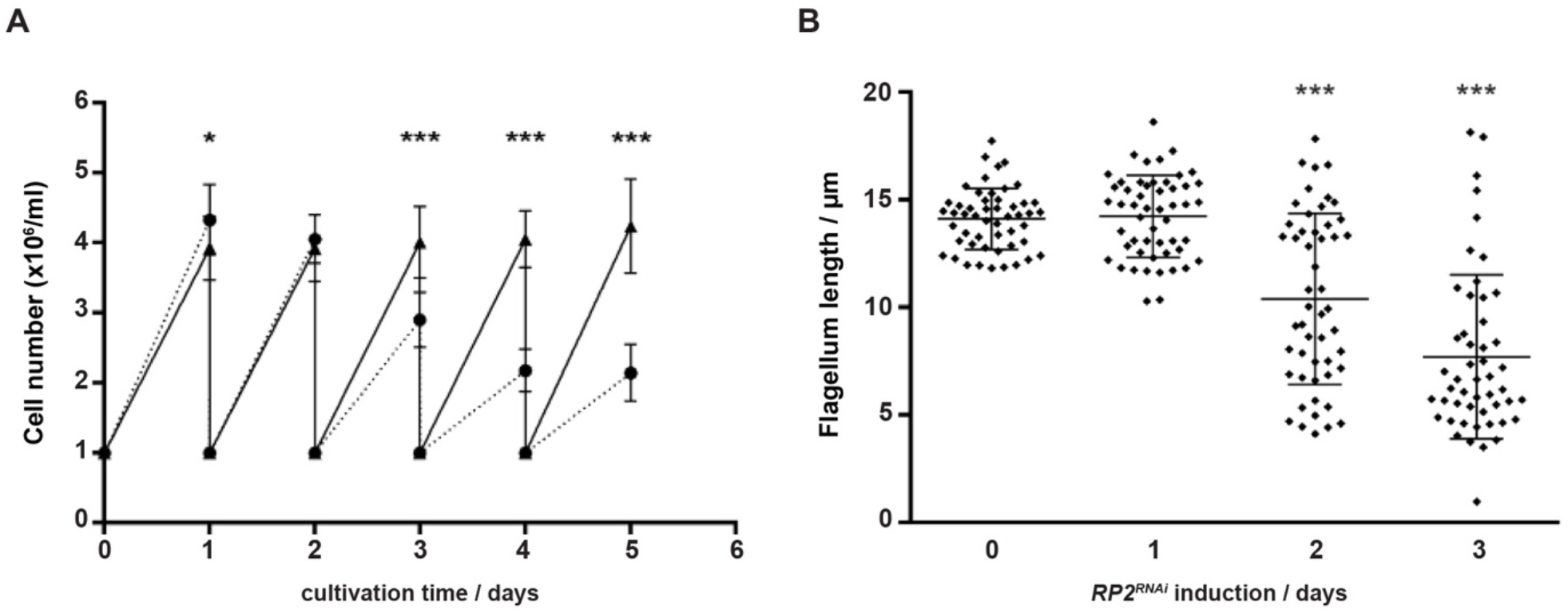
Knockdown of RP2 inhibits cell growth and impacts flagellum length. **(A)** Growth curve of the *RP2*^*RNAi*^ strain with (dashed line and circles) or without tetracycline (plain line and triangles). A severe growth defect emerged on the third day of *RP2* RNAi induction. Similar results were obtained for the *RP2*^*RNAi*^ strain expressing GFP::IFT81 and mCherry::RP2 (data not shown). n ≥ 9 for each time point and condition. **(B)** Flagellum length upon *RP2* RNAi was measured with the mAb25 labelling in methanol-fixed uniflagellated cells. Flagellum length was reduced from the second day of RNAi induction. N ≥ 50 for each condition.

**Figure S3.**
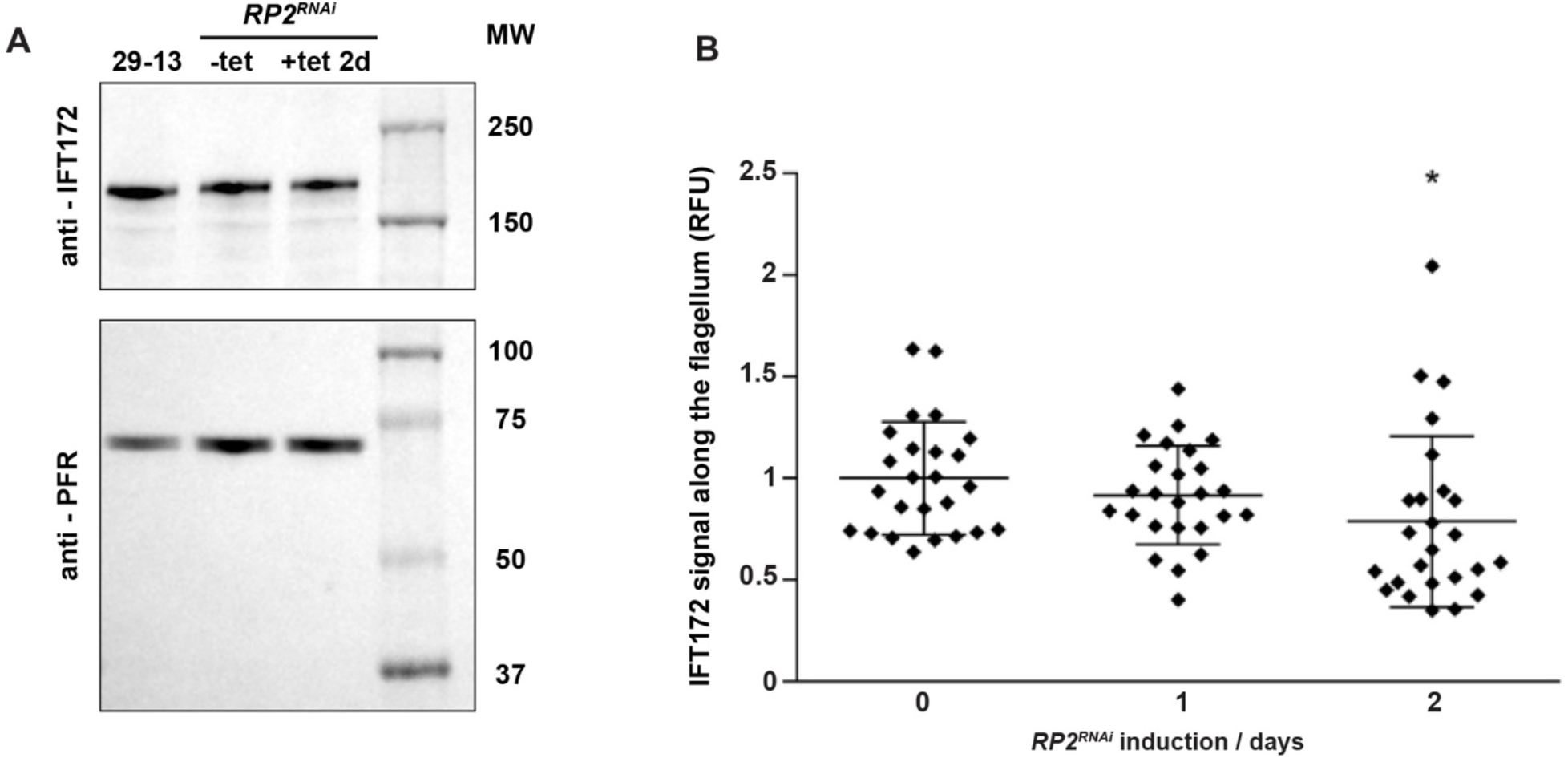
The total amount of IFT172 is not affected in the absence of RP2. **(A)** Western blot analysis of total protein samples from the indicated cell lines probed with either the anti-IFT172 monoclonal antibody (top panel) or the anti-PFR monoclonal antibody L13D6 (bottom panel) as loading control. The position of molecular weight (MW) markers is indicated. **(B)** Quantification of IFT172 signal in the flagellum of uniflagellated cells. Mean IFT172-linked fluorescence was recorded along the flagellum length. Each point stands for an individual cell. n = 25.

**Figure S4.**
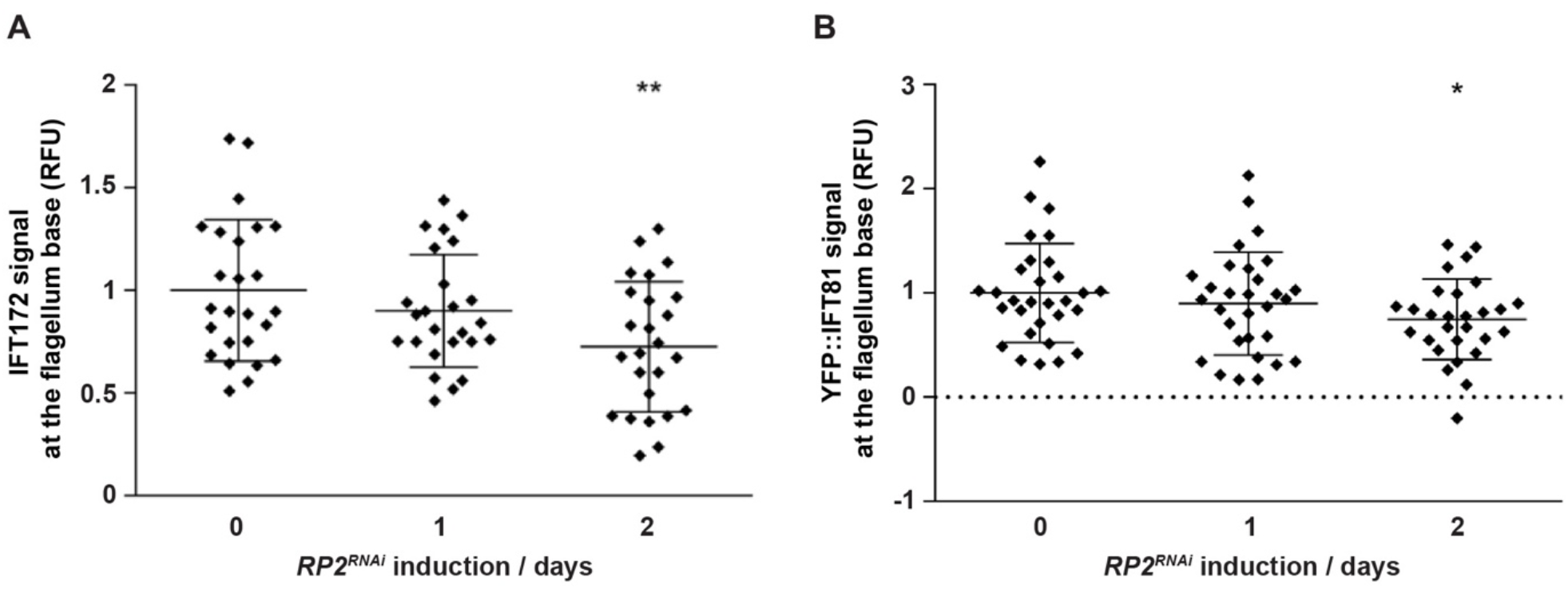
Concentration of IFT proteins at the base of the flagellum is reduced in RP2RNAi induced cells. Fluorescence intensity was measured at the base of the flagellum either following IFA with the anti-IFT172 monoclonal antibody in *RP2*^*RNAi*^ cells **(A)** or using direct YFP fluorescence in live *RP2*^*RNAi*^ cells expressing YFP::IFT81 **(B)**. In both cases, IFT concentration at the base is reduced after two days in RNAi conditions. n ≥ 28 (9-10 parasites from 3 independent experiments).

**Video S1**. FRAP analysis of a non-induced *RP2*^*RNAi*^ cell expressing mNG::IFT81. IFT is recorded for 20 seconds before bleaching the base of the flagellum and follow-up of recovery was monitored over time.

**Video S2**. FRAP analysis of a two-day induced *RP2*^*RNAi*^ cell expressing mNG::IFT81. IFT is recorded for 20 seconds before bleaching the base of the flagellum and follow-up of recovery was monitored over time.

**Video S3**. Live imaging of a non-induced *RP2*^*RNAi*^ biflagellated cell expressing mNG::IFT81. The new flagellum is visible at the posterior end of the cell (left-hand side of the image). Robust IFT can be detected.

**Video S4**. Live imaging of a one-day induced *RP2*^*RNAi*^ biflagellated cell expressing mNG::IFT81. The new flagellum is visible at the posterior end of the cell (left-hand side of the image). IFT is easily detected but the frequency is reduced and several trains are arrested.

**Video S5**. Live imaging of a two-day induced *RP2*^*RNAi*^ biflagellated cell expressing mNG::IFT81. The new flagellum is visible at the posterior end of the cell (left-hand side of the image). IFT trafficking events are severely reduced and several trains are arrested.

